# A Conserved Electrostatic Membrane-Binding Surface in Synaptotagmin-Like Proteins Revealed Using Molecular Phylogenetic Analysis and Homology Modeling

**DOI:** 10.1101/2023.07.13.548768

**Authors:** Nara L. Chon, Sherleen Tran, Christopher S. Miller, Hai Lin, Jefferson D. Knight

## Abstract

Protein structure prediction has emerged as a core technology for understanding biomolecules and their interactions. Here, we combine homology-based structure prediction with molecular phylogenetic analysis to study the evolution of electrostatic membrane binding among vertebrate synaptotagmin-like proteins (Slps). Slp family proteins play key roles in the membrane trafficking of large dense-core secretory vesicles. Our previous experimental and computational study found that the C2A domain of Slp-4 (also called granuphilin) binds with high affinity to anionic phospholipids in the cytoplasmic leaflet of the plasma membrane through a large positively charged protein surface centered on a cluster of phosphoinositide-binding lysine residues. Because this surface contributes greatly to Slp-4 C2A domain membrane binding, we hypothesized that the net charge on the surface might be evolutionarily conserved. To test this hypothesis, the known C2A sequences of Slp-4 among vertebrates were organized by class (from mammalia to pisces) using molecular phylogenetic analysis. Consensus sequences for each class were then identified and used to generate homology structures, from which Poisson–Boltzmann electrostatic potentials were calculated. For comparison, homology structures and electrostatic potentials were also calculated for the five human Slp protein family members. The results demonstrate that the charge on the membrane-binding surface is highly conserved throughout the evolution of Slp-4, and more highly conserved than many individual residues among the human Slp family paralogs. Such molecular phylogenetic-driven computational analysis can help to describe the evolution of electrostatic interactions between proteins and membranes which are crucial for their function.

**Impact statement:** The interior surface of eukaryotic plasma membranes is negatively charged, and many proteins that bind to it have correspondingly evolved a positively charged face. Here, we use techniques from evolutionary biology and computational biophysics to study the conservation of this positively charged surface in an important protein family. We find that the overall surface charge is highly conserved, more so than individual amino acids, consistent with its important role in electrostatic interaction with the membrane.

## Introduction

Membrane-targeting protein domains play many key roles in biology, particularly within eukaryotic secretory pathways. Distributed among several evolutionarily conserved families including C2, PX, and pleckstrin homology (PH) domains, membrane-targeting protein domains are normally soluble in the cytoplasm and drive binding to membrane surfaces in response to signaling events (Lemmon 2008). Typical biochemical signals that proteins recognize include polyanionic phosphoinositide lipid species and/or Ca^2+^ ions (Lemmon 2008; Kutateladze 2010; Corbalan-Garcia & Gomez-Fernandez 2014). In addition, many membrane-binding protein domains also interact with lipid bilayers via nonspecific interactions such as insertion of hydrophobic sidechains into the membrane interior and electrostatic attraction of lysine and arginine residues to anionic phospholipid headgroups (Lemmon 2008). Electrostatic interactions are arguably the most important type of interaction between peripheral proteins and membranes. They are especially important for protein binding to the plasma membrane inner leaflet, which contains approximately 20% phosphatidylserine (PS), 5% phosphatidylinositol (PI), and 1-2% phosphatidylinositol-(4,5)-bisphosphate (PIP_2_) (Murray & Honig 2002; Voelker 2008). While some interactions with phosphoinositides are highly specific, nonspecific electrostatic interactions also play key roles in membrane targeting and affinity. Despite a broad acknowledgment of their significance, systematic quantification of the importance of nonspecific electrostatics in protein-membrane interactions has not been extensively explored. An approach that leverages the dual lens of evolutionary biology and structural biophysics allows for graphical and quantitative interpretation of how natural selection has acted on these protein surfaces. Using this approach can build on prior experimental and computational studies to improve understanding of peripheral protein-membrane interactions.

Here, we use C2A domains from the synaptotagmin-like protein (Slp) family as a model system to explore the evolutionary conservation of the membrane-binding face, which includes a PIP_2_-binding motif in the center of a large positively charged surface (**Figure 1A**). Discovered in the early 2000s, the Slp family consists of five members conserved throughout vertebrates (Slp-1 through Slp-5); at least one is also present in invertebrates (M. Fukuda & Mikoshiba 2001; Serano & Rubin 2003; M. Fukuda 2013). The Slp family is structurally related to synaptotagmins but lack synaptotagmins’ Ca^2+^-sensing role in membrane fusion (Wang et al. 1999; M. Fukuda & Mikoshiba 2001; Bhalla et al. 2008; M. Fukuda 2013). Rather, Slp proteins are Rab27 effectors that function to dock large dense-core secretory vesicles to the plasma membrane prior to exocytosis in a variety of cell types (T. S. Kuroda et al. 2002; M. Fukuda 2013; Izumi 2021). Because Slp family proteins can inhibit exocytosis prior to Ca^2+^ entry, understanding their mechanisms of membrane interaction is important for understanding secretory pathways and their dysfunction in diseases such as diabetes (Tomas et al. 2008; Tsuboi 2009; Izumi 2011). All Slp proteins contain an N-terminal Slp homology domain (SHD) capable of binding Rab GTPases on secretory granules (Mitsunori Fukuda et al. 2001; Chavas et al. 2008), as well as two C-terminal C2 domains, termed C2A and C2B.

**Figure 1:**
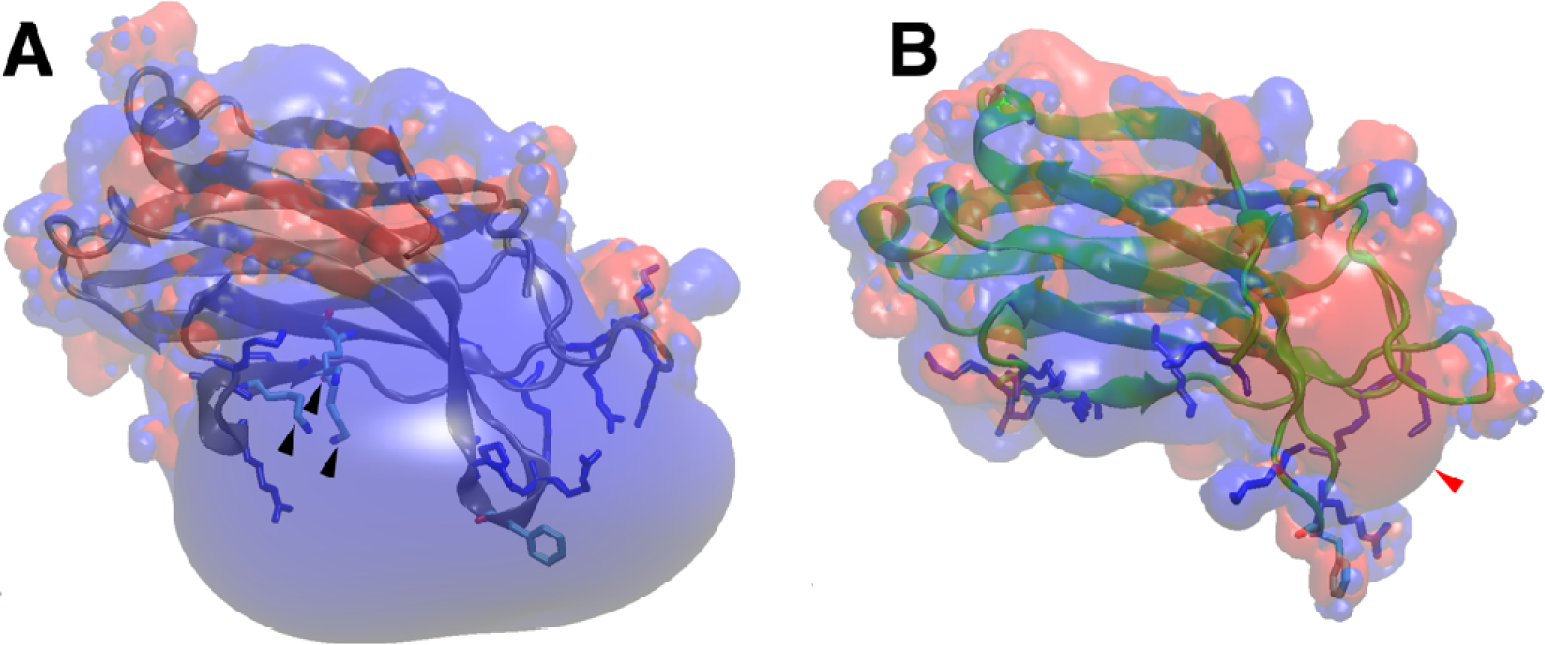
C2 domain structures and electrostatics. **A:** Structure (blue ribbon) and electrostatic surface of the Slp-4 C2A domain (PDB: 3FDW). Black arrowheads indicate the PIP_2_-binding lysine cluster. Other basic residues on the positively charged surface are shown in blue sticks, and the membrane-inserting phenylalanine is shown in cyan. **B:** For comparison, the structure (green ribbon) and electrostatic surface are shown of the synaptotagmin-1 C2A domain in the Ca^2+^-free state (PDB: 1BYN) (Shao et al. 1998). Red arrowhead points to the Ca^2+^-binding pocket, basic residues on the lower surface are shown as blue sticks, and the conserved phenylalanine on calcium-binding loop 3 is shown in cyan. For both structures, surface potentials of +3.5 eV (blue) and -3.5 eV (red) were calculated at 0.15 M ionic strength and pH 7.0.

Originally named from the second conserved domain in protein kinase C (PKC), C2 domains are the 2^nd^-most abundant family of conserved membrane-binding protein domains and are found in over 120 mammalian proteins (Coussens et al. 1986; Nalefski & Falke 1996; Cho & Stahelin 2006; Corbalan-Garcia & Gomez-Fernandez 2014). Two circular permutations of the conserved C2 domain β-strand structure exist in mammalian proteins, termed “Topology I” and “Topology II” (Nalefski & Falke 1996). Several secretory pathway proteins contain two or more C2 domains, including synaptotagmins, Slp proteins, double-C2 (Doc2) proteins, rabphilins, Munc13, ferlins, RIM proteins, and others (Cho & Stahelin 2006; Corbalan-Garcia & Gomez-Fernandez 2014; Dominguez et al. 2022). Some well-known C2 domains, such as those from PKCα and synaptotagmin-1, bind anionic membranes in response to Ca^2+^ influx (Brose et al. 1992; Newton & Keranen 1994; Sutton et al. 1995; Kohout et al. 2002). These C2 domains, which can be Topology I or Topology II, bind one to three Ca^2+^ ions through a conserved set of five anionic residues, typically aspartates (Nalefski & Falke 1996; Murray & Honig 2002). Others lack a full complement of these aspartates but bind membranes containing anionic lipids even in the absence of Ca^2+^ (Yi et al. 2002). A consensus sequence for binding the plasma membrane lipid PIP_2_, including a cluster of three lysine residues centered on a polybasic region in the β_4_ strand, has been recognized among some C2 domains, including both Ca^2+^-dependent and Ca^2+^-independent proteins (Evans et al. 2006; Landgraf et al. 2008; Guerrero-Valero et al. 2009; Lai et al. 2010; Corbalan-Garcia & Gomez-Fernandez 2014).

The role of electrostatics in C2 domain membrane binding has been a topic of interest since the earliest C2 domain structures were reported. It was noted in the 1990s that the PKC-βII C2 domain has several basic residues distributed over a surface rather than in a single strand (Edwards & Newton 1997; Sutton & Sprang 1998). More recently, the presence of a single aspartate residue in the polybasic sequence on the β_4_ strand of the synaptotagmin-1 C2A domain has been shown to cause the weak PIP_2_ selectivity of this domain (Guillen et al. 2013). Although the polybasic region of the synaptotagmin-1 C2A domain has minimal net positive charge due to this aspartate (**Figure 1B**), the surface nevertheless plays a key role in neurotransmitter release (Wu et al. 2022). Thus, the role(s) of nonspecific electrostatic membrane interactions in C2 domain protein function is a longstanding and important question.

Slp family proteins have tandem C2 domains with the Topology I fold that contain the PIP_2_-binding consensus sequence and typically do not require Ca^2+^ for binding membranes (Galvez-Santisteban et al. 2012; Alnaas et al. 2021). [We note, however, that Ca^2+^ is reported to enhance membrane binding of the Slp-3 and Slp-5 C2A domains (M. Fukuda 2002; Taruho S. Kuroda et al. 2002) and to inhibit Slp-2 C2 domains (Yu et al. 2007); the underlying mechanisms have not been fully described.] Thus, they represent a model system for studying electrostatically driven C2 domain membrane binding independent of Ca^2+^. The best-studied Slp family member, Slp-4 (also known as granuphilin), is a key factor in insulin secretion (Izumi et al. 2007; Yamaoka et al. 2015). Previous studies from our lab and others showed that the C2A domain of human Slp-4 binds physiological membranes with high affinity in the absence of Ca^2+^ through nearly equal contributions of (i) the conserved PIP_2_-binding lysine cluster and (ii) a large positively charged surface that includes residues both within the polybasic β_4_ strand and elsewhere on the membrane binding surface (Yu et al. 2007; Lyakhova & Knight 2014; Wan et al. 2015; Alnaas et al. 2021) (**Figure 1**). Mutation of individual residues on the large positively charged surface has little effect on membrane binding affinity, suggesting the attraction is mainly electrostatic and nonspecific; it is also strong enough to drive membrane binding even in the absence of phosphoinositides (Alnaas et al. 2021). In light of this mechanism involving both PIP_2_-selective binding and strong nonspecific binding of background anionic lipids, we set out to explore the evolutionary conservation of the positive charges on this surface.

The goal of the current study is to determine the level of conservation of the large positively charged surface on the C2A domain of Slp-4 and the broader Slp family. We hypothesized that the surface surrounding the PIP_2_-binding pocket evolved to bind membranes containing anionic background lipids with high affinity but low specificity; if so, then the overall charge of the surface might be more highly conserved than individual basic residues. Thus, we compiled the known sequences of C2A domains from vertebrate Slp-4 proteins and the other human Slp family members, predicted structures from consensus sequences, and compared conservation both at the sequence level and among the calculated electrostatic surface potentials.

## Results

### Strategy

The C2A domain of human Slp-4 is the only Slp family C2 domain with a published experimental structure (PDB: 3FDW). Like many C2 domains, its PIP_2_-binding consensus sequence is centered on a triad of lysine residues in the β_3_ and β_4_ strands (Evans et al. 2006; Landgraf et al. 2008; Guerrero-Valero et al. 2009; Lai et al. 2010; Corbalan-Garcia & Gomez-Fernandez 2014; Wan et al. 2015). The broad positively charged surface surrounding this site includes residues from throughout the Slp-4 C2A domain sequence (Alnaas et al. 2021). In order to test the extent of conservation in this nonspecific electrostatic surface, we performed a phylogenetic alignment of Slp-4 C2A domains across vertebrates, extracted consensus sequences, and used homology modeling to construct structures. From these structures, we calculated electrostatic surface maps to visualize the charge distribution on the protein surface. Because the vertebrate Slp-4 C2A domains showed a near-total conservation of both the individual membrane-binding residues and the positively charged surface, we also compared electrostatic surface potentials among homology model structures of C2A domains from the five human Slp family members.

### Conservation in Slp-4 primary structure

Amino acid sequence comparisons reveal a high conservation of membrane-binding residues on Slp-4 C2A domains. Using the Uniref90 database, we identified 218 Slp-4 C2A domain sequences (Uniref90 cluster representatives) with at least 50% identity to human Slp-4, spanning the subphylum vertebrata from fish to mammals. Sequences in the database with <50% identity tended to align more closely with human Slp-5 than Slp-4; therefore, we only analyzed sequences with > 50% identity to human Slp-4. A phylogeny of these sequences shows that the sequences cluster by taxonomic class, as expected (**Figure 2A**), with clear clusters for mammals, birds, amphibians, reptiles, and fishes. The predicted net charges on these protein domain sequences tend to decrease as the level of identity with the human sequence decreases, with the lowest average net charge among the class pisces (**Figure 2B; Table S1**). Interestingly, the trend in net positive charge tends to correlate with the number of lysine residues, while the average count of arginine residues remains relatively constant among the different classes (**Table S1**). Overall, all of the Slp-4 C2A sequences maintain a predicted positive net charge.

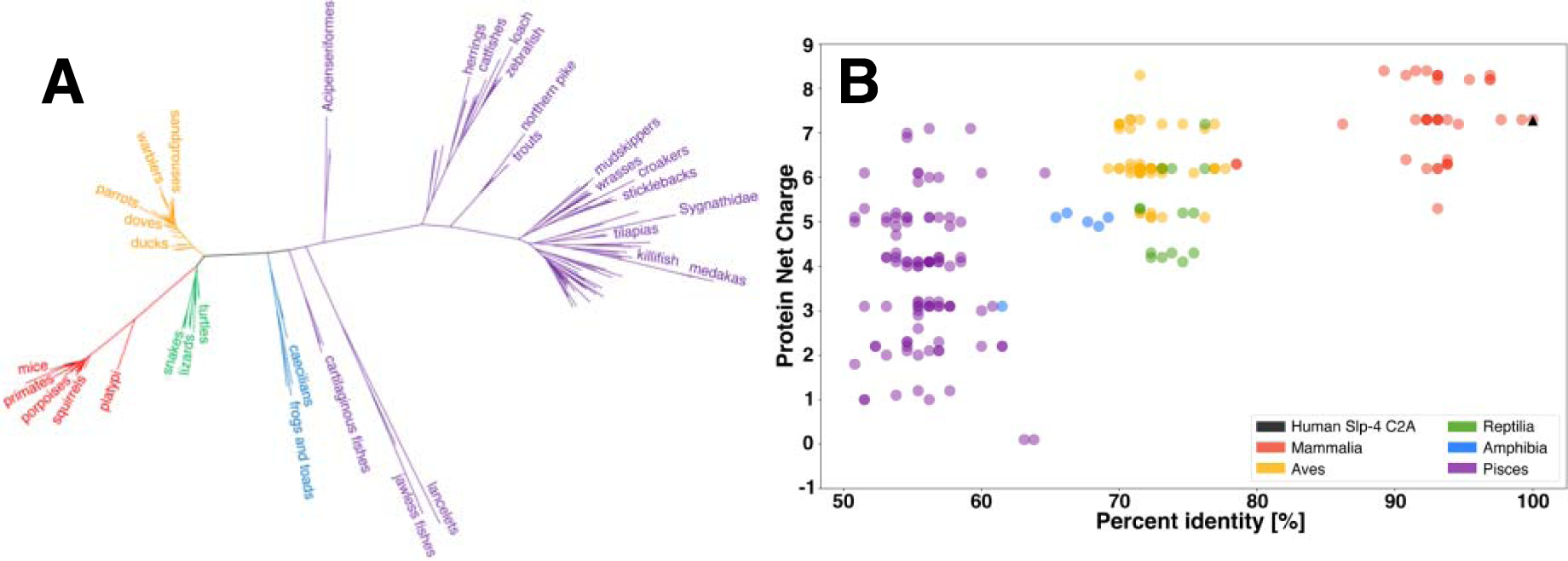

From each taxonomic class (mammalia, aves, reptilia, amphibia, and pisces), a consensus sequence was generated as described in Methods. An alignment of these consensus sequences is shown in **Figure 3**. The alignment shows complete conservation of the consensus PIP_2_ binding site found on many C2 domains, (Corbalan-Garcia & Gomez-Fernandez 2014) which consists of three lysine residues (Lys398, Lys410, and Lys412) along with nearby Tyr396, Trp447, and Asn455 on human Slp-4 (**Figure 3**, red boxes). (For consistency, we report all numbering based on the human Slp-4 sequence.)

**Figure 3:**
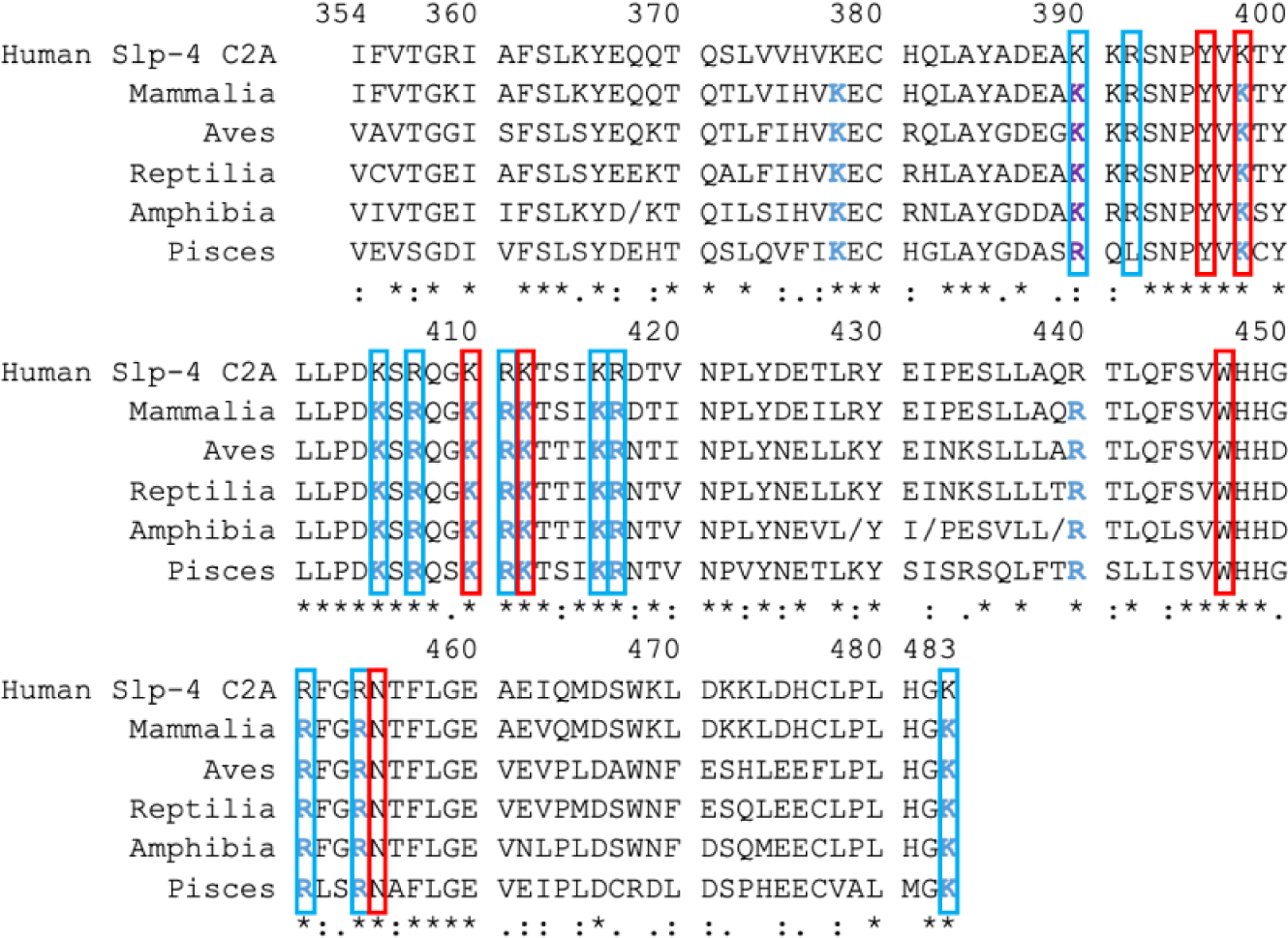
Slp-4 C2A comparison of primary consensus sequences by taxonomic class. Red boxes indicate the consensus sequence for PIP_2_ binding (Guerrero-Valero et al. 2009; Corbalan-Garcia & Gomez-Fernandez 2014), which is completely conserved. Residues in blue boxes are additional ones that made significant contact with anionic lipids in our molecular dynamics simulations of human Slp-4 C2A (Alnaas et al. 2021). Blue text shows residues conserved as K or R; purple text shows residues conserved as basic. Symbols underneath the sequences illustrate the overall level of conservation for each residue: completely conserved (*), strongly similar (:), weakly similar (.). In Amphibia, forward-slashes (/) indicate positions at which two or more residues were equally common among the six sequence clusters: at position 368, Gly (2), His (2) and Ala (2) were found equally; at position 429, Lys (3) and Gln (3); at position 432, Val (3) and Ile (3); at position 439, Val (2) and Ser (2).

Beyond the PIP_2_-binding pocket, our prior study identified ten additional basic residues on the human Slp-4 C2A domain that participate in nonspecific membrane binding and one hydrophobic sidechain that inserts into the nonpolar membrane interior based on molecular dynamics simulations and mutagenesis experiments (Alnaas et al. 2021). The ten basic residues include five in the polybasic β_4_strand region (approximately residues 405-417) and five elsewhere (**Figure 3**). All of these basic residues maintain conserved charge among the consensus sequences except that Arg392 in human Slp-4 corresponds to a leucine in the pisces consensus sequence, although we note that 33% of the pisces sequences contain Lys or Arg at this position. Similarly, the membrane-inserting residue Phe452 is conserved in all consensus sequences except pisces where it is the slightly less hydrophobic leucine. Thus, the membrane-binding residues of the Slp-4 C2A domain are well conserved at the sequence level.

### Electrostatic surface maps show high conservation in Slp-4

The electrostatic surface of the Slp-4 C2A domain is highly conserved from fish to humans. We threaded each of the consensus sequences in **Figure 3** onto the published human Slp-4 C2A crystal structure using SwissModel (Waterhouse et al. 2018) and calculated electrostatic surfaces using APBS (**Figure 4**) (Baker et al. 2001). Strikingly, the positively charged membrane-binding surface (shown as a blue surface extending in the -z direction) is highly similar among all the classes of the protein. The volume of the positively charged surface is slightly smaller in the pisces consensus structure, corresponding to the presence of leucine at position 392 instead of arginine. The charged surface on the region near Phe452 in the mammalian structure is greatly positive. However, some negative charge is visible in that region in the other classes. These small areas stem in part from the presence of an Asp at position 450 in most of the non-mammalian structures.

**Figure 4.**
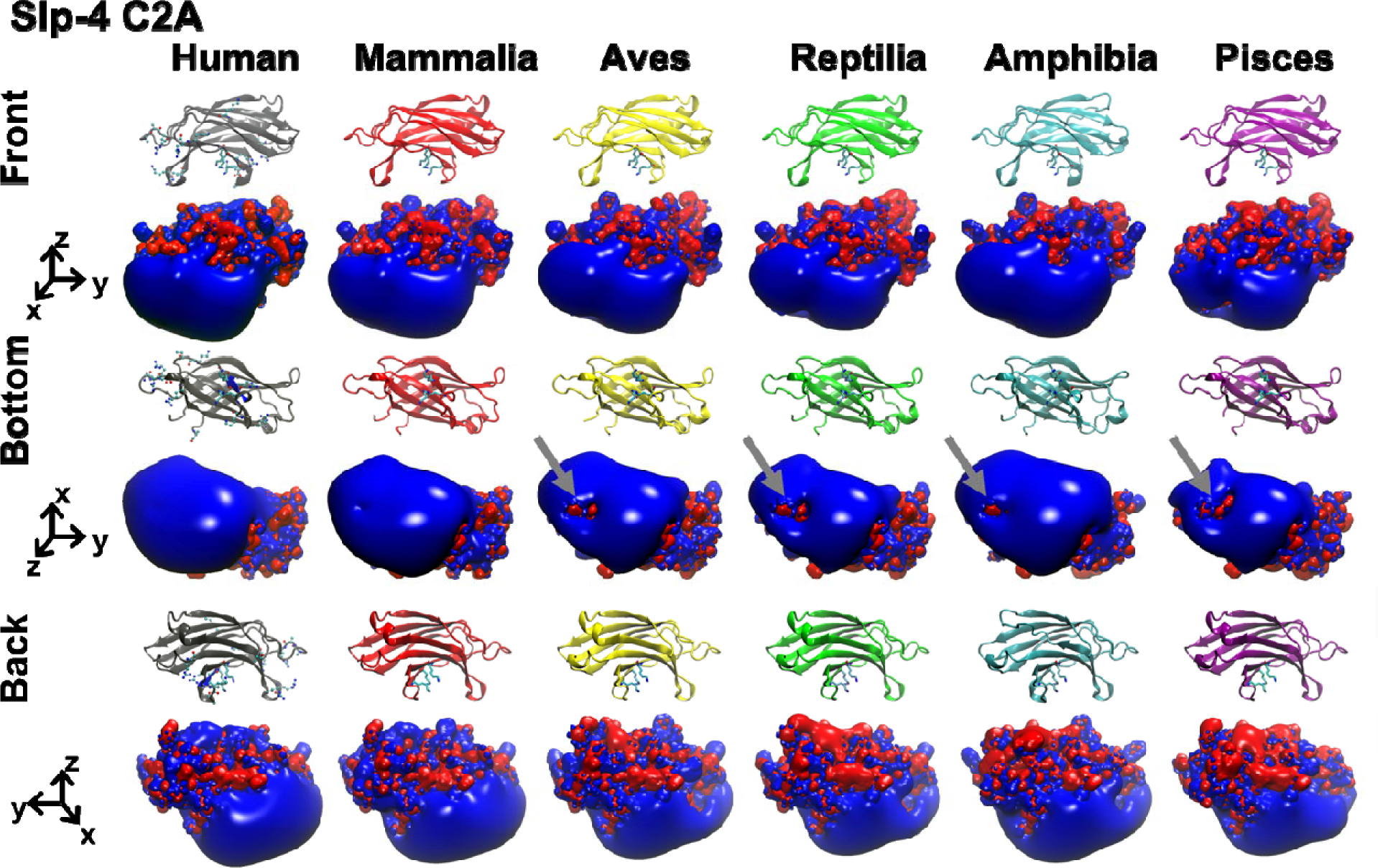
Electrostatic surfaces of Slp-4 C2A domain consensus structures. The homology model structures and electrostatic surface charge maps are shown for the five consensus sequences of Mammalia (red), Aves (yellow), Reptilia (green), Amphibia (blue), and Pisces (purple) with the crystal structure of the human Slp-4 C2A (gray). In the electrostatic surfaces, red represents negative charge potential (-3.5 eV) while blue represents positive charge potential (+3.5 eV). Gray arrows indicate partial negative charge near the hydrophobic Phe452.

These differences in the positively charged membrane binding surface are much smaller than the electrostatic differences in the surface pointing away from the membrane (+z direction in **Figure 4**), which vary considerably among the different classes. Also at the primary sequence level, conservation in the +z surface is lower than on the membrane-facing surface (**Figure S1**). The function of this +z surface is unknown. Clearly, the membrane-facing surface of this domain is highly conserved in both sequence and electrostatics.

In order to quantify the similarity in the positively charged surface, we used the webPIPSA tool which was developed for comparing electrostatic protein-protein interaction surfaces (Richter et al. 2008). We calculated similarity indices (SI) for the human and each class consensus structure versus every other class consensus structure using several different volumes within the membrane-facing surface as regions of comparison (**Figure 5 and Figure S2-S3**). The full SI scale ranges from zero (identical) to 1.4 (uncorrelated) to 2.0 (fully anti-correlated). Focusing on the electrostatics around Phe452, Slp-4 C2A pairs all had SI values of 0.28 or less, indicating strong correlation (**Figure 5**). Correlations were even stronger when comparing the region immediately around the conserved lysine cluster, and slightly less strong when comparing the region around Arg454 (**Figure S3**).

**Figure 5.**
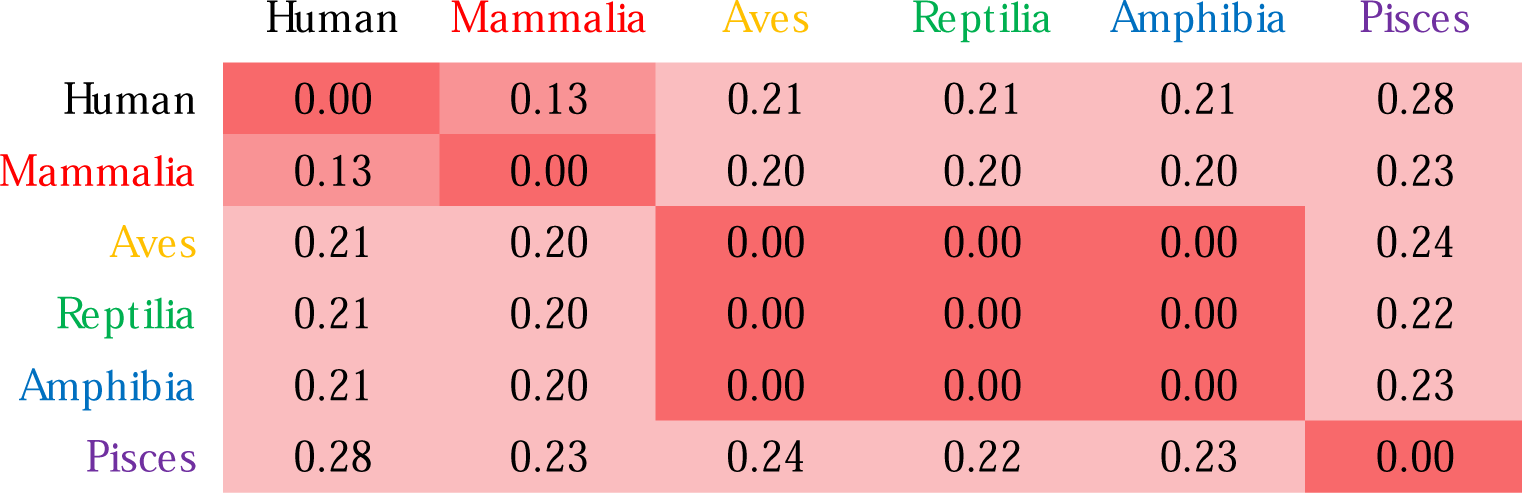
Quantitative comparison of electrostatic surface charge in Slp-4 C2A homology model structures. Comparison values were calculated using webPIPSA for a sphere of 10-Å radius centered on the coordinates of Phe452 Cα in the crystal structure of the human Slp-4 C2A. For comparisons using other regions, see **Figure S3**. Smaller values (red) represent more similar electrostatic surface charge profiles. The matrix is symmetrical.

### Electrostatic surface comparison among human Slp paralogs

The analysis above shows that the Slp-4 C2A membrane-binding surface is highly conserved both in sequence and electrostatics. If the positively charged membrane-binding surface is an evolutionarily conserved feature important for protein function, one prediction is that the overall positive charge of this surface among the C2A domains of the larger family of synaptotagmin-like proteins may be more strongly conserved than individual residues. In order to test this prediction, we expanded our comparison to the sequences and predicted structures of all five human Slp C2A domains.

Some of the membrane-binding residues in Slp-4 C2A are conserved among Slp family proteins while others are not (**Figure 6**). Five of the six consensus residues for PIP_2_ binding are completely conserved, and the sixth (Asn455) is conserved in four of the five proteins. Of the ten additional membrane-binding basic residues we previously identified in Slp-4, five are completely conserved as basic while the other five are not completely conserved. For the non-conserved residues, the proteins that lack a basic residue at that position sometimes have one nearby; for example, Slp-1 and Slp-2 lack an Arg at position 451 but contain an Arg at 449 that is not present in Slp-4.

**Figure 6:**
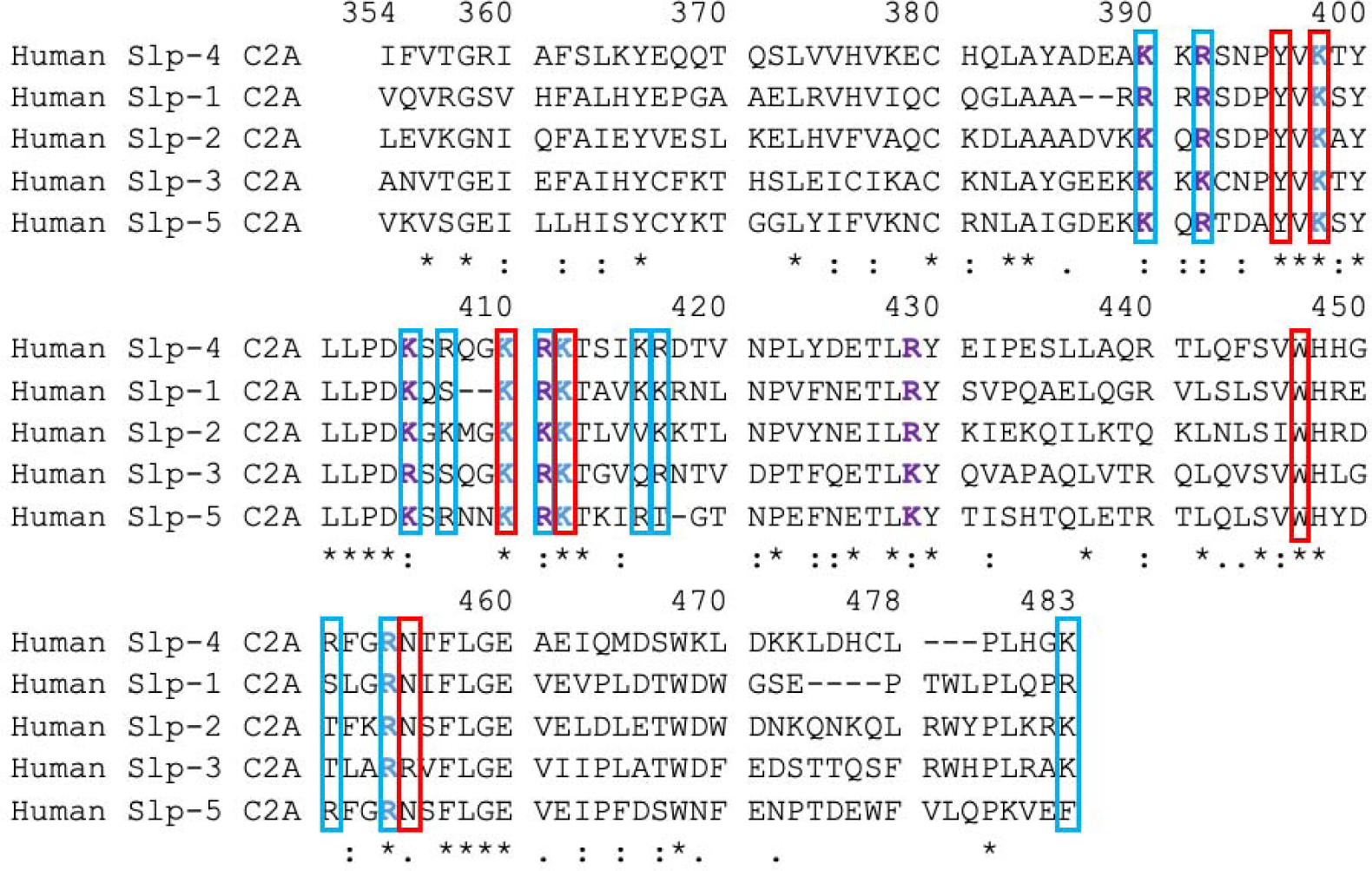
Alignment of human Slp C2A domain sequences. The top line shows the numbering of Slp-4 C2A for consistency in comparisons. Sequence ranges of other proteins shown are 626-758 in Slp-2, 267-391 in Slp-1, 304-436 in Slp-3, and 404-536 in Slp-5. Red boxes indicate the consensus sequence for PIP_2_ binding (Wan et al. 2015), which is completely conserved. Residues in blue boxes are additional ones that made significant contact with anionic lipids in our molecular dynamics simulations of human Slp-4 C2A (Alnaas et al. 2021). Blue text shows residues strictly conserved; purple text shows residues conserved as basic. Symbols underneath the sequences illustrate the overall level of conservation for each residue: completely conserved (*), strongly similar (:), weakly similar (.) as defined by the Gonnet PAM 250 scoring matrix (Gonnet et al. 1992). Dashes indicate that a given residue is absent in the sequence.

Electrostatic surfaces of the homology model structures of human Slp-1 through Slp-5 show differences in the size and location of the positively charged surface, consistent with the sequence comparison (**Figure 7**). The positive lobe is largest in Slp-1 and Slp-2, is shifted toward the -y direction in Slp-1, and is smaller and shifted more toward the +y direction in Slp-5. However, strong positive charge is clearly present on the -z surface of all five protein domains, whereas the electrostatics of the +z surfaces vary substantially among the five family members. Quantitative comparison using webPIPSA confirms that the similarities are relatively strong, with most SI values less than 0.5 (**Figure 8 and Figure S4**). Although the similarity patterns differ depending on the precise region of comparison, the regions near the conserved lysine cluster in particular show very high similarity (**Figure S4**). Together, these results support the assertion that electrostatic membrane binding is an evolutionarily conserved feature of the Slp family C2A domains, as the presence of a large positively charged surface is more strongly conserved than several of the individual basic residues on that surface.

**Figure 7.**
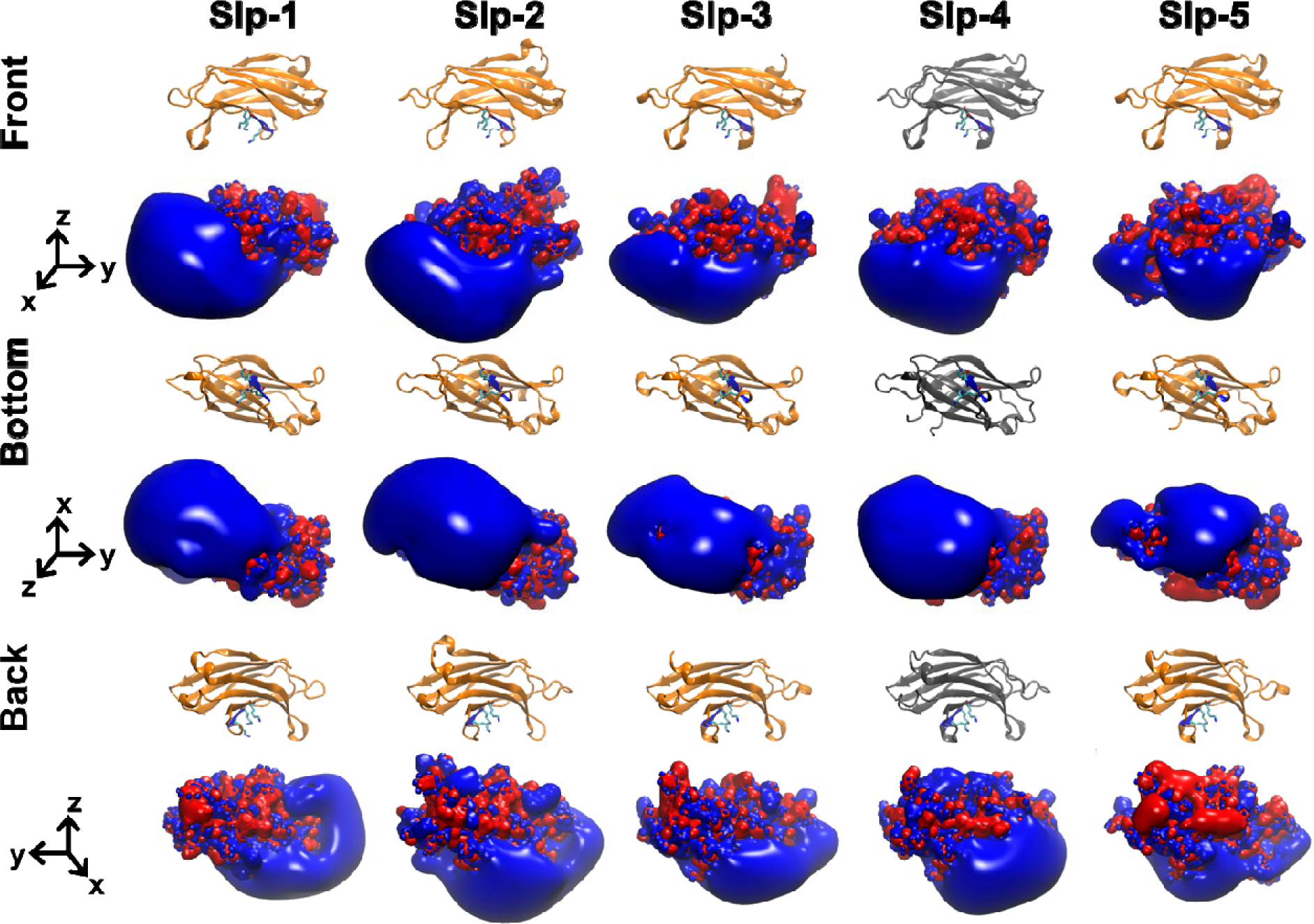
Electrostatic surfaces of human Slp family C2A domains. The electrostatic surface charge distributions are shown for the homology models of the C2A domains from human Slp-1, Slp-2, Slp-3, and Slp-5 along with the crystal structure of the human Slp-4 C2A domain (gray). In the electrostatic surfaces, red represents negative charge potential (-3.5 eV) while blue represents positive charge potential (+3.5 eV).

**Figure 8.**
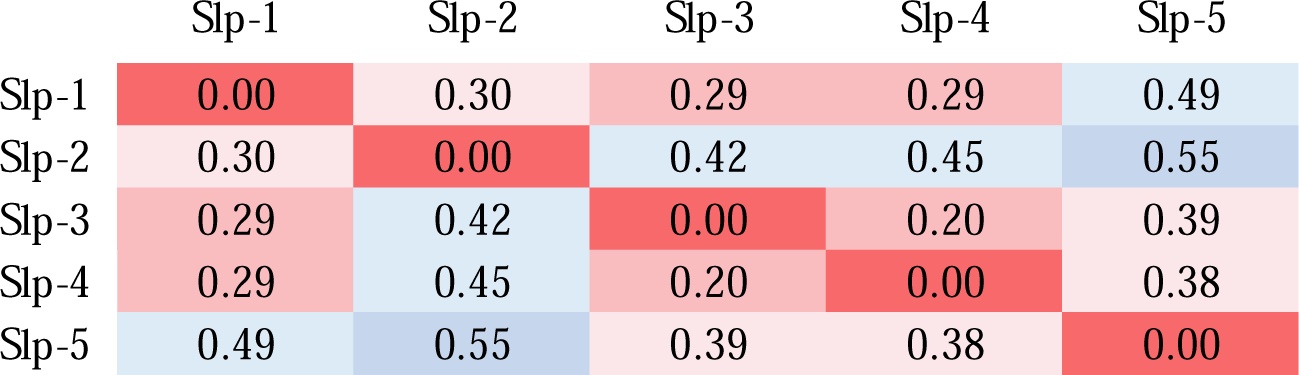
Quantitative comparison of electrostatic surface charge in human Slp family C2A homology model structures. Comparison values were calculated for each model structure using webPIPSA for a sphere of 10-Å radius centered on the coordinates of Phe452 Cα in the human Slp-4 crystal structure. Smaller values (red) represent more similar electrostatic surface charge profiles. For comparisons using other regions, see **Figure S4**.

## Discussion

### Importance of electrostatic surfaces

Proteins interact with lipid membranes through a combination of nonspecific electrostatic attractions, the hydrophobic effect, and specific molecular recognition. In particular, electrostatic attraction can contribute greatly toward overall binding affinity for a negatively charged membrane surface. In biological systems, both the affinity and the specificity of interactions are tuned to match the native environment. Coincidence detection is a common feature of peripheral protein-lipid interactions (Medkova & Cho 1999; Johnson et al. 2000; Corbin et al. 2004; Evans et al. 2006; Lemmon 2008; Ziemba et al. 2014), which allows for membrane specificity and affinity to be tuned via separate sites or regions of a protein. In vertebrates, nonspecific electrostatic attractions contribute not only to affinity through Coulombic attraction, but also to membrane specificity because the plasma membrane inner leaflet is enriched in anionic lipids relative to internal membranes such as ER and Golgi (Corbin et al. 2007; van Meer et al. 2008). Thus, investigating evolutionary conservation can offer clues to the importance of electrostatics in protein-lipid interactions.

In this study, we have used phylogenetic analysis and structure prediction to compare the evolutionary conservation of a positively charged membrane-binding surface from C2A domains of Slp family proteins. These bind membranes through (1) a conserved PIP_2_ binding site; (2) a large positively charged surface surrounding the PIP_2_ binding site; and (3) a single membrane-inserting hydrophobic residue. It is clear from simple sequence analysis that the PIP_2_-binding residues are highly conserved (**Figures 3 and 6**). Our electrostatic analysis further shows that the large positively charged surface is also a conserved feature of this family of proteins. Notably, the results indicate the overall surface charge on the membrane binding surface is more highly conserved than several of the individual basic residues outside of the PIP_2_ binding site. Not surprisingly, individual basic residues are more highly conserved among vertebrate orthologs of Slp-4 than when comparing across human Slp family paralogs.

### Sequence conservation among Slp proteins and relatives

Slp family proteins are closely evolutionarily related to synaptotagmins, rabphilins, and Doc2b proteins, all of which contain tandem C2 domains with Type I topology (M. Fukuda & Mikoshiba 2001; Rickman et al. 2004; Craxton 2010). Many but not all synaptotagmins as well as rabphilins and Doc2b C2 domains bind membranes in a Ca^2+^-dependent manner (Pinheiro et al. 2016). The Ca^2+^-dependent synaptotagmin C2 domains typically contain five conserved Asp residues in a Ca^2+^-binding pocket: two in the β_2_-β_3_ loop and three in the β_6_-β_7_ loop (**Figure 9**). However, Slp family C2A domains lack most or all of these Asp residues: vertebrate Slp-4 consensus sequences contain one (mammalia, pisces) or two (aves, reptilia, amphibia) Asp residues at the calcium-binding positions, and the rest of the human Slp domains contain between zero (Slp-3) and three (Slp-2 and Slp-5) (**Figure 9**). This pattern is consistent with a functional loss of Ca^2+^ binding by Slp C2A domains after diverging from a common ancestor with synaptotagmins and other related proteins. Interestingly, Slp C2A domains contain a conserved His at one of these positions (H448 in human Slp-4) whose role is unknown. Its position at the base of the pocket suggests it could function as a pH sensor analogous to some PH domains (He et al. 2008).

**Figure 9.**
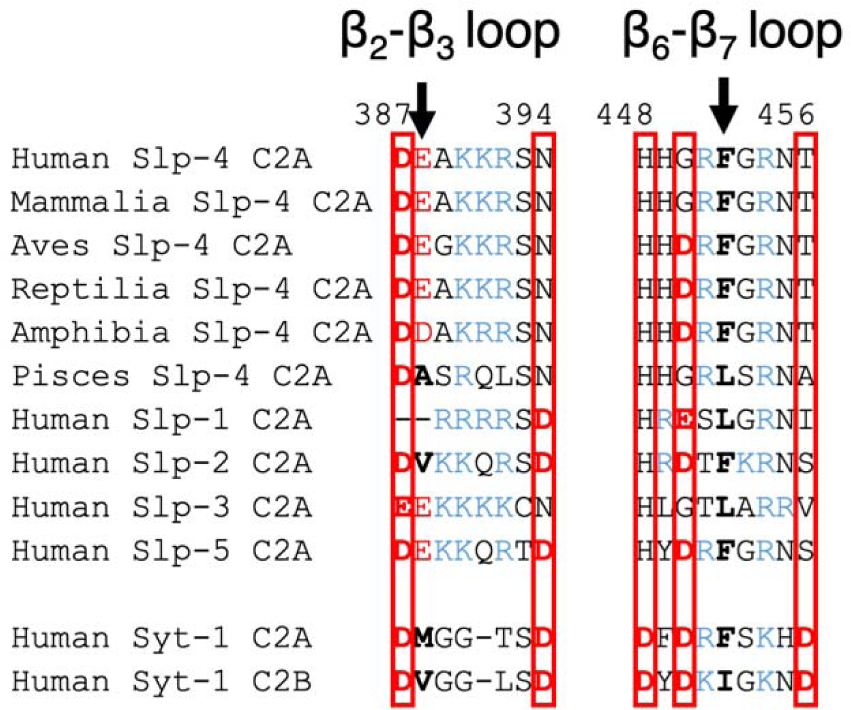
Slp C2A comparison to Ca^2+^-dependent C2 domains of synaptotagmin-1 (Syt-1). The β_2_-β_3_ and β_6_-β_7_ loop sequences are shown (also called CBL1 and CBL3, respectively, in Ca^2+^-dependent C2 domains). Asp and Glu residues are shown in red; Lys and Arg are shown in blue. Boxes indicate positions of conserved Asp in Ca^2+^-binding C2 domains. Arrows indicate positions of conserved hydrophobic character in synaptotagmins; nonpolar residues at these positions are in bold.

The Slp-4 C2A domain also contains a single hydrophobic residue at the tip of its β_6_-β_7_ loop, corresponding to Phe452 in human Slp-4, which inserts into the membrane interior during molecular dynamics simulations (Alnaas et al. 2021). This residue was Phe or Leu in all sequences analyzed in **Figure 3 and 6**. In contrast, C2A domains of membrane-binding synaptotagmins typically have a hydrophobic residue at this position plus another one on the β_2_-β_3_ loop (**Figure 9**) (X. Zhang et al. 1998; Osterberg et al. 2015). Among the Slp sequences analyzed, only human Slp-2 (Val) and pisces Slp-4 (Ala) had hydrophobic residues at this second position, suggesting that membrane insertion of the β_2_-β_3_ loop may be another feature that is being lost following divergence from the last common ancestor with synaptotagmins. In Slp C2A domains, the Phe or Leu on the β_6_-β_7_ loop is surrounded by a large lobe of positive charge (**Figures 4 and 7**), although the presence of Asp or Glu residues add small patches of negative charge nearby in some Slp-4 sequences (**Figure 4**) and human Slp-5 (**Figure 7**).

The conserved motif by which Slp C2A domains bind PIP_2_ is present in many C2B domains of synaptotagmins, but not the C2A domains of Ca^2+^-dependent synaptotagmins (Guillen et al. 2013; Alnaas et al. 2021). To our knowledge, no evolutionary analysis of synaptotagmin electrostatic surfaces has been conducted, although electrostatic surface maps have been reported for some individual synaptotagmin C2 domains (Chon et al. 2015; MacDougall et al. 2018). It would be interesting to compare the level of conservation seen in the Slp C2A positively charged surface to those of other tandem C2 domain proteins.

### Usefulness of this approach to studying protein-membrane interactions

To our knowledge, this is the first study to combine evolutionary and structural modeling to study conservation of a membrane-binding electrostatic surface among a family of C2 domain proteins. Typical prior approaches address C2 domain evolutionary relationships at the sequence level to determine key consensus motifs and/or identify novel C2 domain relatives beyond the conventional classification (Craxton 2010; D. Zhang & Aravind 2010; Farrell et al. 2012; D. Zhang & Aravind 2012; Tellez-Arreola et al. 2022). Several studies have observed that Ca^2+^ switches the electrostatic profile of many C2 domains from negative to positive, beginning with the seminal work of Murray and Honig (Murray & Honig 2002). A more recent study has performed detailed molecular dynamics simulations of selected Ca^2+^-independent C2 domains, showing broad binding to anionic lipids including PIP_2_ (Larsen & Sansom 2021). Other recent work has taken advantage of structure prediction software to clarify C2 domain folds and identify hidden domains (Dominguez et al. 2022). The approach we describe here is well suited to explore the evolutionary conservation of an electrostatic protein surface and thereby demonstrate its functional importance. These broad electrostatic surface interactions are key for binding to anionic membranes, but they are beyond the scope of co-evolution methods that have been developed for identifying evolutionarily conserved protein-protein interactions from sequence analysis (Lockless & Ranganathan 1999), or more recent learning approaches that use (in part) underlying evolutionary information to infer molecular interactions stabilizing protein complexes (Humphreys et al. 2021; Bryant et al. 2022). In general, we note that methods for learning protein-membrane interactions that integrate physical models and large-scale data sets seem to lag behind related methods used to infer protein-protein interactions.

This study represents a preliminary demonstration of the approach of investigating conserved electrostatic membrane-binding surfaces by using the Slp protein family as a starting example. While our approach combined automated sequence alignment and homology modeling with manual curation of sequences, future work could leverage recent advances in structure prediction (Jumper et al. 2021) to pursue broader comparisons of membrane binding protein families and could conceivably be extended to unrelated proteins whose surfaces have converged on an electrostatically driven membrane binding function. In addition, future work could combine evolutionary analyses of electrostatic and hydrophobic surfaces, as both modes of nonspecific binding work together to determine the affinity of peripheral membrane proteins for their target membranes. For Slp family proteins, the electrostatic attraction of the conserved positively charged surface appears to be a major, evolutionarily conserved driver of target membrane affinity.

## Methods

### Slp-4 sequence search and protein net charge calculations

The human Slp-4 C2A sequence was used to search the Uniref90 database to obtain an initial selection of 500 clustered sets of sequences using the MMseqs2 algorithm (Steinegger & Söding 2018). Any given clustered sequences with <50% sequence identity to the human Slp-4 C2A domain and/or containing uncharacterized proteins, Slp-4 alternative splicing isoforms, or other Slp family members were removed from the initial selection. To refine the sequence sets, a molecular phylogenetic tree was built and clades were manually curated with taxonomic information. Any sequences that produced unbranched lineages outside the clades were subjected to deletion. As a result, a total of 218 representative sequences from UniRef90 were retained for further analysis. Of the 218 Slp-4 sequences, 38 were class Mammalia, 60 were Aves, 14 were Reptilia, 6 were Amphibia, and 100 were Pisces. A FASTA file of these sequences along with the human reference sequence are included in the **Supporting Information**, along with a spreadsheet containing sequence ID numbers and further annotation of each sequence. A breakdown of the percent identity of each sequence versus the human Slp-4 C2A sequence is shown in **Figure S5**.

A multiple sequence alignment of the retained Slp-4 C2A domains was built using COBALT (Papadopoulos & Agarwala 2007) based on conserved domain and local sequence similarity information among the sequences. The gap penalties were set as follows: gap opening penalty of -11, gap extension penalty of -1, end gap opening penalty of -5, and end gap extension penalty of -1. C2A domains were manually curated after the multiple sequence alignment by comparison to the human sequence in the published crystal structure (PDB: 3FDW). A new phylogenetic tree for the selected Slp-4 C2A domains was constructed using the Maximum Likelihood method in MEGA-X (version 10.1.8) with the Jones-Taylor-Thornton substitution matrix (Jones et al. 1992; Kumar et al. 2018). The Nearest-Neighbor-Interchange heuristic search (Li et al. 1996) was used for topological transformation, and the tree was bootstrapped 100 times. The tree was visualized in iToL (Letunic & Bork 2021).

### Consensus sequences per taxonomic class

The consensus sequence for each vertebrate taxonomic class was determined by identifying the most abundant residue at each position. All consensus sequences were built and aligned using the AlignIO module in the Bio.Align Python package (Cock et al. 2009). The consensus sequences had the following percent identity when compared to the human Slp-4 C2A: 95% for Mammalia, 72% for Aves, 77% for Reptilia, 69% for Amphibia, and 61% for Pisces.

### 3D Homology Models and Electrostatic Surface Charge Calculations

The crystal structure of the human Slp-4 C2A (PDB ID: 3FDW.pdb) was used to construct the 3D homology models for the five consensus sequences of the Slp-4 C2A vertebrate classes through SWISS-MODEL (Waterhouse et al. 2018). Then the electrostatic surface charges were calculated by utilizing the webserver of APBS (Baker et al. 2001). All calculations were done at pH 7.0 and 0.15 M KCl. Although homology modeling has limitations at low sequence identities, the high level of homology among Slp-4 C2A consensus sequences (all ≥ 61% identity to the human Slp-4 C2A template) inspires good confidence in the inferred structures, including the large positively charged membrane binding surfaces.

Electrostatic surface charges were quantified and compared with webPIPSA (Protein Interaction Property Similarity Analysis) (Richter et al. 2008) at 300 K and 0.15 M ionic strength. For the data shown in **Figures 5 and 8**, analysis of the membrane-facing region centered on the coordinates of the alpha carbon of Phe452 in the human Slp-4 C2A crystal structure (3FDW), with a 10-Å radius from this point defining the volume of interest in each of the computed structures. Other regions of analysis are described in Figure S2 for data shown in Figures S3 and S4. The electrostatic potential distances for each model were computed on a one-to-one basis. The distance was judged by the similarity index (SI) as computed by webPIPSA: 0 for identical, 1.4 for fully uncorrelated, and 2.0 for fully anti-correlated.

### Sequence alignment of the human Slp family C2A domains

The human C2A domain sequences for Slp-1 through Slp-5 obtained through BLASTP were aligned using the Clustal Omega web service in UniProt. The similarity among residues at each position was determined by the Gonnet Point Accepted Mutation (PAM) 250 scoring matrix (Gonnet et al. 1992). Homology models and electrostatic surface charges were calculated as described above.

## Conflicts of Interest

The authors declare no conflicts of interest.

## Supplementary Material

*Chon_etal_2023_Supporting Information.pdf*: PDF file containing Figures S1-S6 and Table S1.

*Chon_etal_2023_all_Slp4_sequences.fasta*: FASTA file listing all Slp-4 sequence clusters analyzed.

*Chon_etal_2023_Supplemental_Sequence_data.xlsx*: Spreadsheet file listing all Slp-4 sequence clusters analyzed along with sequence ID number and other annotations.

*Chon_etal_2023_SwissModel_structures.zip*: Compressed file containing 9 homology-generated protein structure PDB files and the human Slp-4 C2A reference structure.

## Supporting information

Supplement: Structures

Supplement: Sequence Alignment

Supplement: FASTA sequences

## Acknowledgments

This work was supported by NIH grant R15GM102866. Additional support provided by the Camille and Henry Dreyfus Foundation (TH-14-028 to H.L. and TH-18-061 to J.K.) and by the CU Denver Undergraduate Research Opportunity Program to S.T.

## Notes

### Competing Interest Statement

The authors have declared no competing interest.

### Summary of Updates

Introduction updated to clarify nomenclature of C2 domains and historical significance of positively charged surfaces on C2 domains; Figure 1 added; webPIPSA analysis revised to include more regions of comparison; all sections revised for clarity.

