## Supplemental Tables and Figures for "A Conserved Electrostatic Membrane-Binding Surface in Synaptotagmin-Like Proteins Revealed Using Molecular Phylogenetic Analysis and Homology Modeling"

**Table S1.** Slp-4 C2A per-class data of percent identities (vs. human), the formal net charge at pH 7.0, and the average number of particular charged residues in each class. Values are given as mean +/- S.D. For comparison, the human Slp-4 C2A domain has a net charge of +7.3.

| | % Identity | Net Charge | Average Counts | | | | |
|---|---|---|---|---|---|---|---|
| | | | Lys | Arg | His | Asp | Glu |
| Mammalia | 93 ± 4 | 7.3 ± 0.8 | 13.8 ± 1.0 | 8.3 ± 1.0 | 6.1 ± 0.6 | 7.1 ± 0.4 | 8.0 ± 0.5 |
| Aves | 72 ± 2 | 6.3 ± 0.6 | 11.4 ± 0.8 | 8.7 ± 0.8 | 4.9 ± 0.7 | 4.1 ± 0.6 | 9.8 ± 0.5 |
| Reptilia | 74 ± 2 | 5.2 ± 1.0 | 11.7 ± 0.6 | 8.4 ± 0.7 | 5.5 ± 0.9 | 4.4 ± 1.2 | 10.7 ± 0.9 |
| Amphibia | 66 ± 3 | 4.7 ± 0.8 | 10.2 ± 0.8 | 9.0 ± 1.1 | 3.7 ± 1.0 | 7.3 ± 0.5 | 7.2 ± 0.4 |
| Pisces | 56 ± 3 | 3.8 ± 1.6 | 9.8 ± 1.7 | 8.8 ± 1.6 | 4.3 ± 1.2 | 7.4 ± 1.4 | 7.5 ± 1.2 |

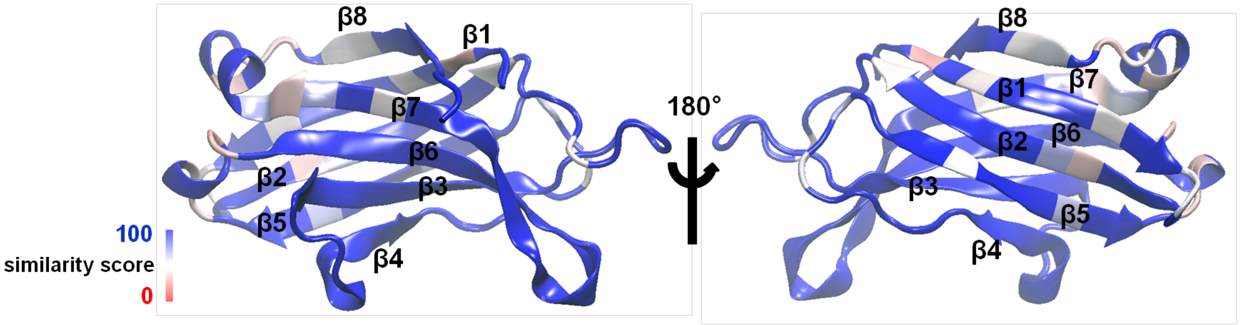


**Figure S1. Slp-4 membrane binding face is highly conserved.** Shown is a sequence similarity comparison between human Slp-4 C2A (PDB: 3FDW) and an overall consensus sequence using the most frequent residue at each position from a multiple sequence alignment using all species. Interestingly, the membrane binding surface (β3-4 and β6) had ~100 % identity (blue) to the human Slp-4 C2A.

**Human Slp-4 C2A**
**F452 CA (7.081, -17.021, 62.684)**
***r* = 10.0 Å**

**Human Slp-4 C2A**
**R454 CZ (-0.154, -19.644, 61.122)**
***r* = 10.0 Å**

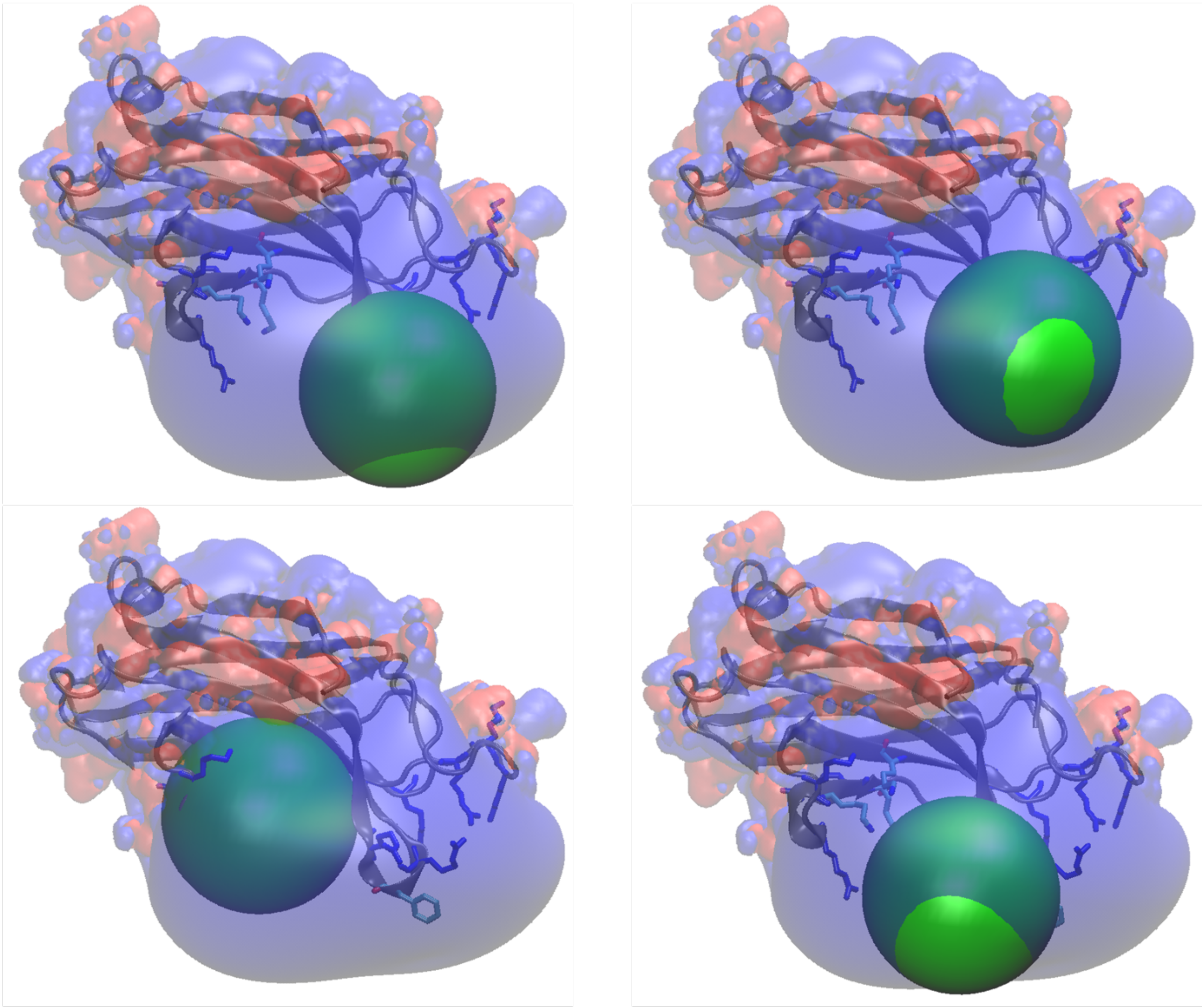

**Human Slp-4 C2A**
**Lysine Cluster (4.161, -8.903, 49.108)**
***r* = 10.0 Å**

**Human Slp-4 C2A**
**Offset (4.320, -20.562, 55.861)**
***r* = 10.0 Å**

**Figure S2. Regions used for comparison of electrostatics via webPIPSA.** The region around Phe452 Cα was used for webPIPSA results in the main manuscript, and the other three regions were used for data in Figures S3 and S4. The green spheres illustrate the regions used for each analysis, and the numbers above or below each structure show the radius and coordinates of each center point. PDB files are available in the Supporting Information.

**A** Slp-4 C2A R454 CZ

| | Human | Mammalia | Aves | Reptilia | Amphibia | Pisces |
|---|---|---|---|---|---|---|
| Human | 0.00 | 0.06 | 0.44 | 0.35 | 0.37 | 0.21 |
| Mammalia | 0.06 | 0.00 | 0.46 | 0.38 | 0.40 | 0.24 |
| Aves | 0.44 | 0.46 | 0.00 | 0.13 | 0.10 | 0.25 |
| Reptilia | 0.35 | 0.38 | 0.13 | 0.00 | 0.04 | 0.18 |
| Amphibia | 0.37 | 0.40 | 0.10 | 0.04 | 0.00 | 0.20 |
| Pisces | 0.21 | 0.24 | 0.25 | 0.18 | 0.20 | 0.00 |

**B** Slp-4 C2A Lysine cluster

| | Human | Mammalia | Aves | Reptilia | Amphibia | Pisces |
|---|---|---|---|---|---|---|
| Human | 0.00 | 0.09 | 0.09 | 0.10 | 0.08 | 0.08 |
| Mammalia | 0.09 | 0.00 | 0.06 | 0.06 | 0.06 | 0.04 |
| Aves | 0.09 | 0.06 | 0.00 | 0.00 | 0.00 | 0.04 |
| Reptilia | 0.10 | 0.06 | 0.00 | 0.00 | 0.04 | 0.04 |
| Amphibia | 0.08 | 0.06 | 0.00 | 0.04 | 0.00 | 0.04 |
| Pisces | 0.08 | 0.04 | 0.04 | 0.04 | 0.04 | 0.00 |

| Color Bar |
|---|
| 0.0 |
| 0.1 |
| 0.2 |
| 0.3 |
| 0.4 |

**C** Slp-4 C2A Offset

| | Human | Mammalia | Aves | Reptilia | Amphibia | Pisces |
|---|---|---|---|---|---|---|
| Human | 0.00 | 0.08 | 0.15 | 0.14 | 0.13 | 0.09 |
| Mammalia | 0.08 | 0.00 | 0.19 | 0.17 | 0.17 | 0.11 |
| Aves | 0.15 | 0.19 | 0.00 | 0.00 | 0.04 | 0.09 |
| Reptilia | 0.14 | 0.17 | 0.00 | 0.00 | 0.00 | 0.08 |
| Amphibia | 0.13 | 0.17 | 0.04 | 0.00 | 0.00 | 0.06 |
| Pisces | 0.09 | 0.11 | 0.09 | 0.08 | 0.06 | 0.00 |

**Figure S3. Similarity Indices from webPIPSA calculations of alternative protein regions across Slp-4 C2A evolution.** Calculations were performed as described in Methods, except using the regions of interest shown in Figure S2.

**A** Human Slp C2A R454CZ

| | Slp-1 | Slp-2 | Slp-3 | Slp-4 | Slp-5 |
|---|---|---|---|---|---|
| Slp-1 | 0.00 | 0.26 | 0.48 | 0.66 | 0.27 |
| Slp-2 | 0.26 | 0.00 | 0.56 | 0.74 | 0.41 |
| Slp-3 | 0.48 | 0.56 | 0.00 | 0.36 | 0.55 |
| Slp-4 | 0.66 | 0.74 | 0.36 | 0.00 | 0.69 |
| Slp-5 | 0.27 | 0.41 | 0.55 | 0.69 | 0.00 |

**B** Human Slp C2A Lysine cluster

| | Slp-1 | Slp-2 | Slp-3 | Slp-4 | Slp-5 |
|---|---|---|---|---|---|
| Slp-1 | 0.00 | 0.36 | 0.26 | 0.31 | 0.21 |
| Slp-2 | 0.36 | 0.00 | 0.33 | 0.26 | 0.35 |
| Slp-3 | 0.26 | 0.33 | 0.00 | 0.18 | 0.28 |
| Slp-4 | 0.31 | 0.26 | 0.18 | 0.00 | 0.24 |
| Slp-5 | 0.21 | 0.35 | 0.28 | 0.24 | 0.00 |

**C** Human Slp C2A Offset

| | Slp-1 | Slp-2 | Slp-3 | Slp-4 | Slp-5 |
|---|---|---|---|---|---|
| Slp-1 | 0.00 | 0.35 | 0.38 | 0.31 | 0.17 |
| Slp-2 | 0.35 | 0.00 | 0.36 | 0.32 | 0.38 |
| Slp-3 | 0.38 | 0.36 | 0.00 | 0.17 | 0.43 |
| Slp-4 | 0.31 | 0.32 | 0.17 | 0.00 | 0.33 |
| Slp-5 | 0.17 | 0.38 | 0.43 | 0.33 | 0.00 |

Color Bar: 0.0 0.1 0.2 0.3 0.4 0.5 0.6 0.7

**Figure S4. Similarity Indices from webPIPSA calculations of alternative protein regions across the human Slp family.** Calculations were performed as described in Methods, except using the regions of interest shown in Figure S2.

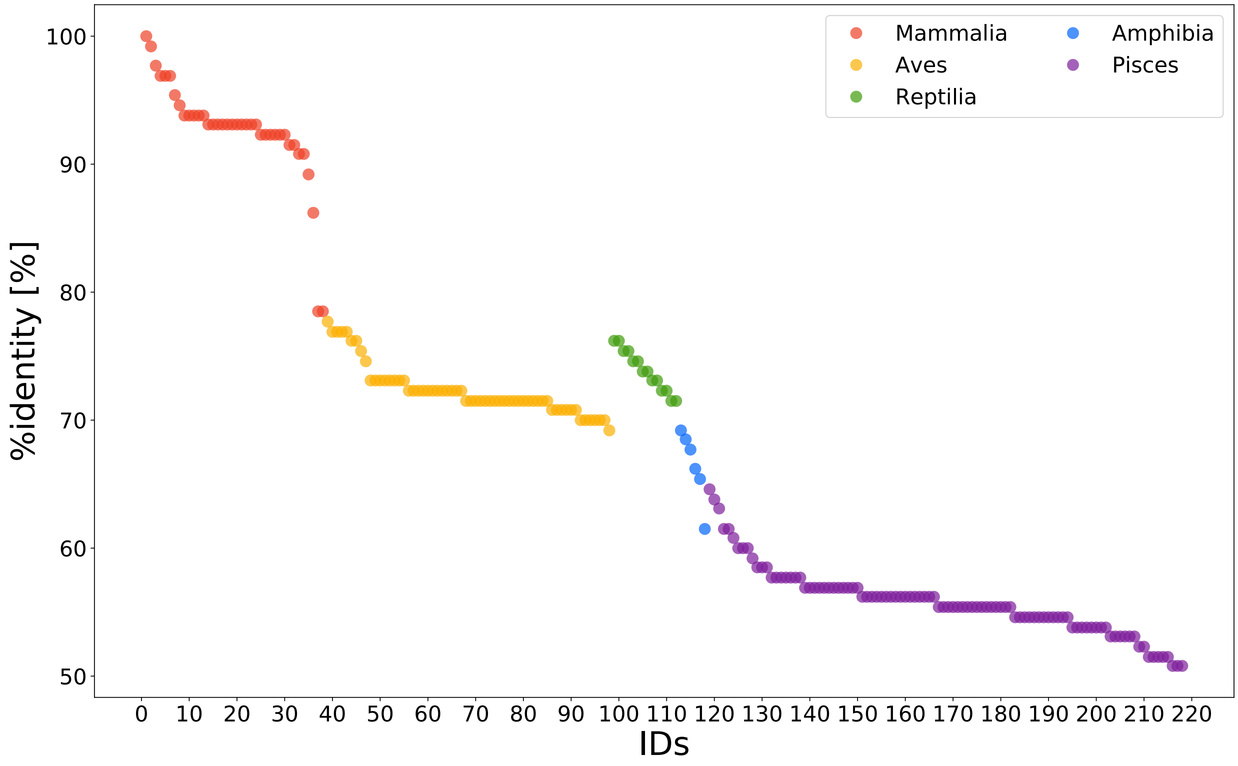


**Figure S5.** Sequence identity comparison among Slp-4 C2A domains. IDs are sorted by taxonomic class (see inset legend) followed by percent identity to the human reference sequence (%identity). Full sequence data including IDs are provided in an Excel spreadsheet in the Supporting Information.

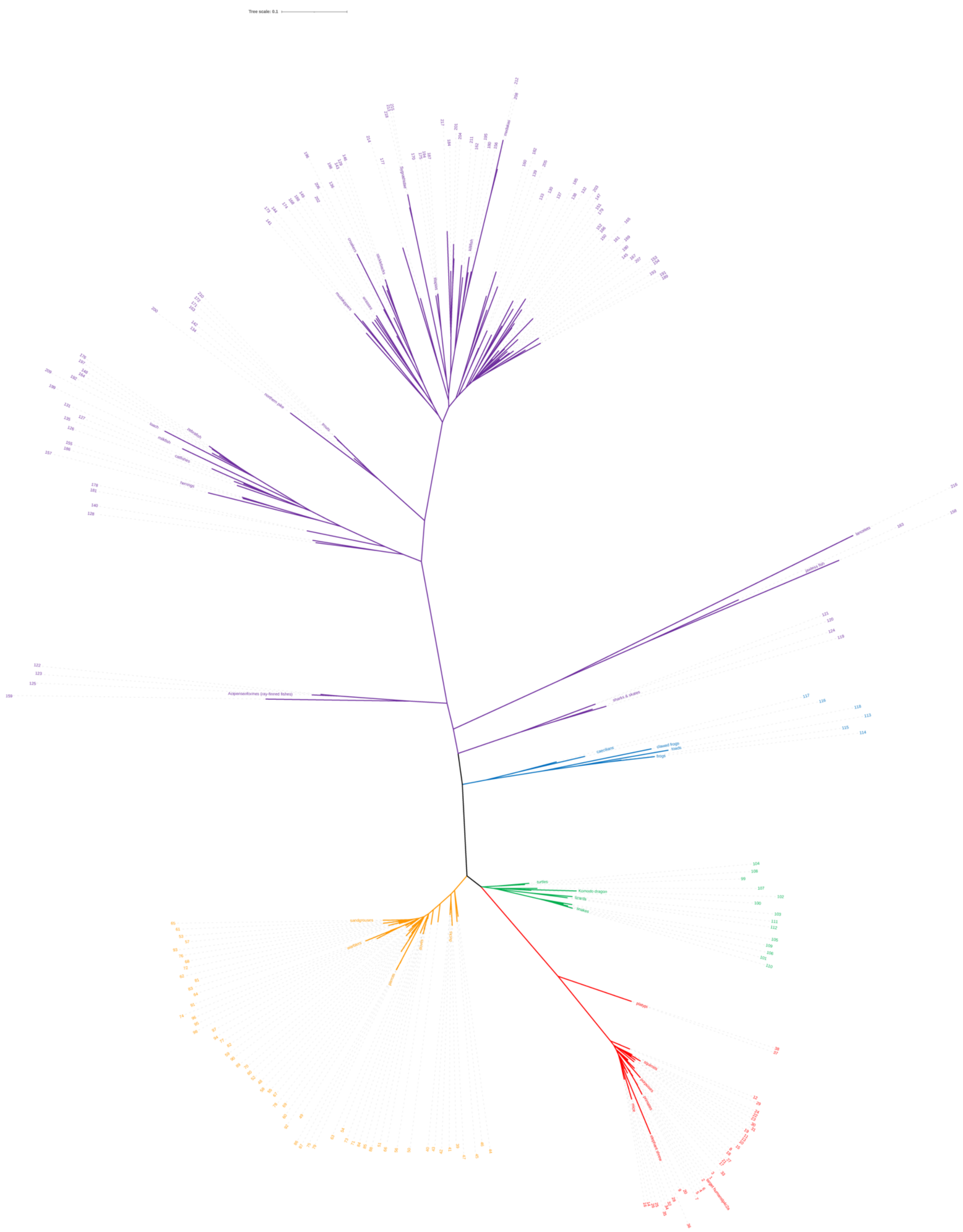

**Figure S6:** Detail of Figure 2A. Numbers on each leaf correspond to the cluster numbers in the Excel file in the Supporting Information.
